# ATR Inhibitors as Potent Modulators of DNA End Resection Capacity

**DOI:** 10.1101/2020.01.13.905059

**Authors:** Diego Dibitetto, Jennie R. Sims, Carolline F.R. Ascenção, Kevin Feng, Raimundo Freire, Marcus B. Smolka

## Abstract

DNA end resection is a key step in homologous recombination-mediated DNA repair. The ability to manipulate resection capacity is expected to be a powerful strategy to rationally modulate DNA repair outcomes in cancer cells and induce selective cell lethality. However, clinically compatible strategies to manipulate resection are not yet well established. Here we find that long-term inhibition of the ATR kinase has a drastic effect on DNA end resection. Inhibition of ATR over multiple cell division cycles depletes the pool of pro-resection factors and prevents RAD51 as well as RAD52-mediated DNA repair, leading to toxic end-joining and hypersensitivity to PARP inhibitors. The effect is markedly distinct from acute ATR inhibition, which blocks RAD51-mediated repair but not resection and RAD52-mediated repair. Our findings reveal a key pro-resection function for ATR and define how ATR inhibitors can be used for effective manipulation of DNA end resection capacity and DNA repair outcomes in cancer cells.

## Introduction

DNA replication is a major source of DNA double-strand breaks (DSBs), which arise as replication forks encounter nicks on DNA or collide with obstacles such as DNA-protein or DNA-DNA crosslinks, actively transcribed genes and hard-to-replicate sequences^1^. The ability of cells to sense and repair replication-induced lesions heavily relies on the *ataxia-telangiectasia*-mutated (ATM)-rad3-related kinase ATR^2^. ATR, together with its cofactor ATRIP, is recruited to RPA-coated single-stranded DNA (ssDNA) exposed at replication-induced lesions and DSB intermediates^3^. Upon recruitment, ATR becomes activated by the proteins ETAA1 and TOPBP1 to initiate an extensive signaling response^4–8^. In its canonical mode of action, ATR phosphorylates and activates the CHK1 kinase, which has established roles in the control of cell cycle progression and transcriptional responses, among other processes^9–11^. While chemical or genetic ablation of ATR or CHK1 function results in loss of viability and exquisite sensitivity to replication stress^11–15^, the mechanisms by which these kinases maintain genome integrity are still enigmatic. In particular, it remains unclear how ATR and CHK1 control DNA repair processes necessary to repair DSBs generated during DNA replication.

Recently, ATR has emerged as an important regulator of homologous recombination (HR)^16–18^. HR is initiated by the 5’-3’ nucleolytic processing of DNA ends (referred to as resection), which allows subsequent recruitment of the RAD51 recombinase^19^. Resection initially requires the activity of the MRN (MRE11-RAD50-NBS1) nucleolytic complex together with the stimulatory factors BRCA1 and CTIP^20–24^. Short ssDNA overhangs generated by MRN are then further processed by the concerted activity of the exonuclease EXO1, the flap-endonuclease DNA2, and the helicase BLM^24,25^. ssDNA intermediates generated by resection robustly activate ATR^26–30^, which then controls RAD51 loading by directly phosphorylating PALB2^16^, a tumor suppressor required for the recruitment of BRCA2-RAD51 to resected breaks^31^. In addition, ATR indirectly promotes HR capacity through the activation of E2F transcription and the consequent expression of HR proteins during the S-phase of the cell cycle^17^. Mechanistically, ATR-CHK1 signaling mediates the release of E2F6, E2F7 and E2F8 repressors from target promoters allowing the E2F1 activator to initiate gene transcription^32,33^. However, it remains unclear how the depletion of E2F-regulated HR factors alters the steps of HR and the impacts for DNA repair outcomes.

Most cancer cells exhibit intrinsically high levels of ATR activation, which has been attributed to their increased levels of replication stress caused by oncogene-induced de-regulation of DNA replication^12,17^. Given the higher reliance of cancer cells on ATR signaling, inhibition of ATR signaling has been explored as a strategy for cancer therapy^13,34–36^. Early development of potent ATR inhibitors enabled recent clinical trials for the treatment of different malignancies, including prostate, ovarian, and lung cancer^37,38^. Numerous ATR inhibitors have been developed since then, and a range of clinical trials are currently on phase I and phase II^38,39^. Notably, ATR inhibitors were found to exhibit strong synergism with PARP-inhibitors in sensitizing cancers^17,40–43^, although the mechanism behind such synergy remains elusive.

Here, using chemical and genetic approaches to manipulate ATR signaling, we find that ATR inhibition severely impairs DNA resection, the initial step of homology-directed DNA repair. Long-term treatment with sub-lethal doses of different ATR inhibitors led to a significant depletion of BRCA1, CTIP, and BLM, three essential DNA end resection factors. Loss of these resection factors correlates with a substantial reduction in DNA end resection capacity as measured quantitatively through a CRISPR-Cas9-based assay, and hypersensitivity to PARP inhibitors. Our results support a mechanism by which long-term ATR inhibition is more effective at hyper-sensitizing cells to PARP inhibitors compared to short term ATR inhibition. We find that loss of DNA end resection after prolonged suppression of ATR signaling sensitized cells to PARP inhibitors in a DNA-PKcs-dependent manner and propose that long-term ATR inhibition allows NHEJ-mediated repair and the subsequent accumulation of toxic chromosomal aberrations. Short-term ATR inhibition, while effective at suppressing canonical HR by preventing RAD51 loading, has little impact on resection, and therefore allows engagement of alternative rescuing repair pathways, including RAD52-mediated repair. Overall, our findings reveal a key pro-resection function for ATR and define how ATR inhibitors can be used for effective manipulation of DNA end resection capacity and DNA repair outcomes in cancer cells.

## Results

### Chemical and genetic ablation of ATR signaling depletes the abundance of key resection factors

We have previously shown that long-term treatment with the ATR inhibitor (ATRi) VE-821 severely depletes the abundance of HR factors and reduces HR capacity in cancer cells^17^. Here, using two distinct ATR inhibitors, VE-821, and AZD6738, we find that the abundance of three central resections factors, BRCA1, BLM and, CTIP, is strongly reduced by long-term ATR inhibition in U-2OS cells (Figs. 1a-d). Of importance, only minor alterations in cell cycle distribution were observed under the conditions used (Fig. 1e), indicating that the observed changes in protein abundance are not due to the indirect effects of a cell cycle arrest. To further confirm that the diminished abundance of DNA end resection factors was caused by loss of ATR signaling, we monitored the abundance of these proteins upon genetic ablation of the ATR activators TOPBP1 and ETAA1. We used an HCT116-derivative cell line where the ATR Activating Domain (AAD) in the *ETAA1* gene has been removed by CRISPR-Cas9, and both alleles of *TOPBP1* were tagged with an mAID epitope to conditionally induce TOPBP1 degradation upon auxin treatment^44,45^ (Fig. 1f). TOPBP1 auxin-dependent degradation resulted in destabilized BRCA1, BLM, and CTIP (Fig. 1g), similar to the effect observed with ATRi treatment. The abundance of resection factors was restored after auxin washout, indicating that loss of resection capacity is transient and is caused by the temporary and reversible suppression of ATR signaling (Fig. 1h). Importantly, auxin-induced TOPBP1 depletion did not alter the cell cycle distribution (Fig. 1i). Taken together, these results show that ATR signaling plays a key role in maintaining the abundance of crucial pro-resection factors. Since genotoxins are not used in the described experiments, the findings suggest that the maintenance of resection factor abundance relies on intrinsic ATR activation. Furthermore, since acute treatment (up to 24 hours) with ATR inhibitors does not result in similar depletion of resection factors, the activity of ATR must be inhibited over multiple cell division cycles for the altered abundances to become noticeable.

**Fig. 1.**
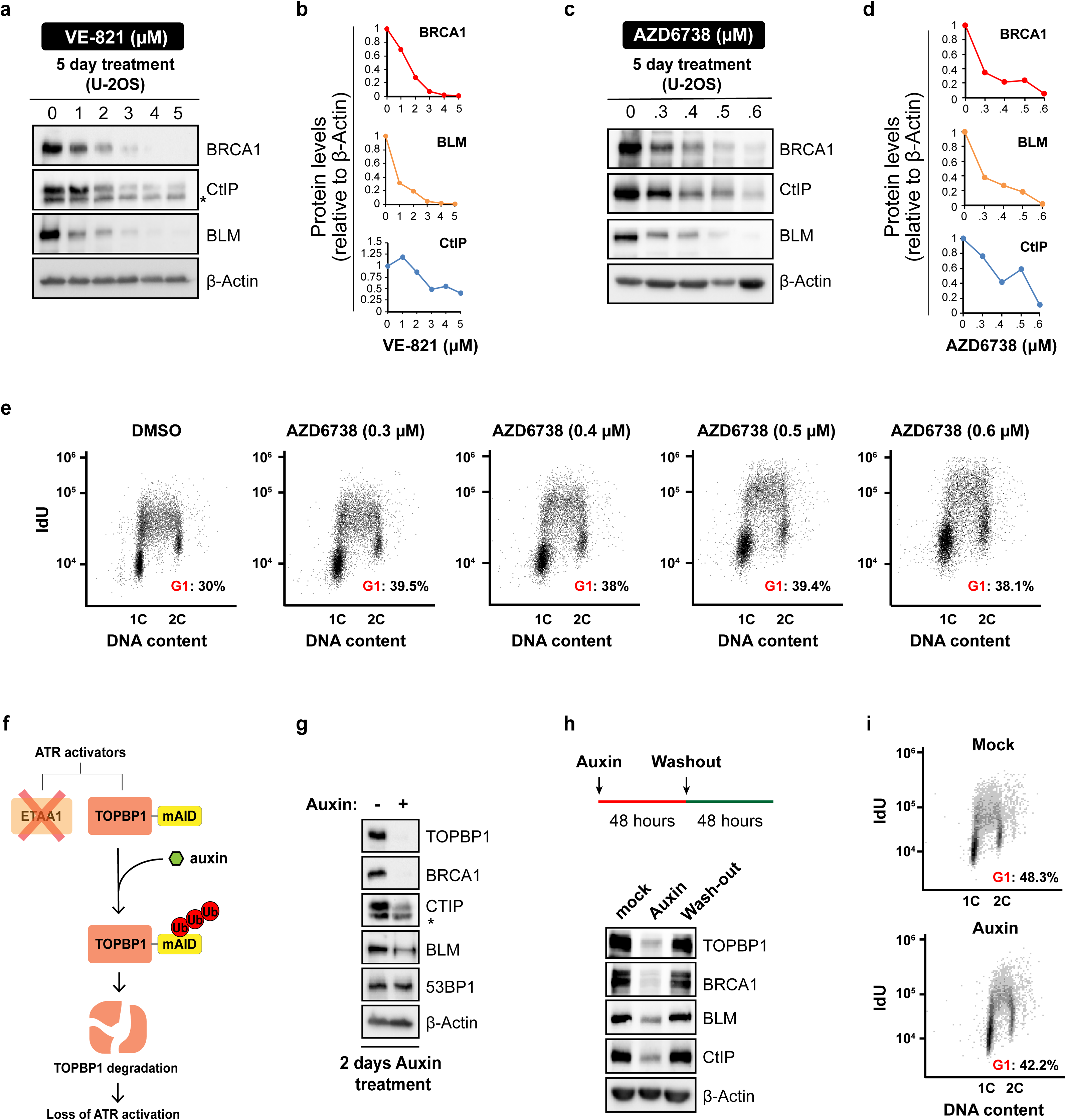
Chemical and genetic ablation of ATR signaling depletes the abundance of key resection factors. (**a**) U-2OS cells were cultured for 5 days in medium containing DMSO or the indicated concentrations of ATR inhibitor (ATRi) VE-821 and analyzed by immunoblotting. (**b**) Quantification of blots in (**a**). (**c**) U-2OS cells were treated as in (**a**) but with the ATRi AZD6738. (**d**) Quantification of blots in (**c**). (**e**) IdU incorporation analysis of U-2OS cells treated as in (**c**). (**f**) Strategy for abrogating ATR activators using the HCT116-*ETAA1ΔAAD*-TOPBP1-mAID cell line. (**g**) Immunoblot analysis in HCT116-*ETAA1ΔAAD*-TOPBP1-mAID cells after 2 days in auxin. (**h**) Immunoblot analysis of HCT116-*ETAA1ΔAAD*-TOPBP1-mAID cells treated for 2 days with auxin and released in fresh medium for additional 2 days. (**i**) IdU incorporation analysis of HCT116-*ETAA1ΔAAD*-TOPBP1-mAID cells treated as in (**g**).

### Long-term ATR inhibition severely impairs DNA end resection

Based on the results above, we predicted that long-term ATR inhibition leads to a strong decrease in DNA end resection efficiency. To test this, we used an engineered system to introduce DSBs at a defined genomic locus through CRISPR-Cas9 technology and measured nearby ssDNA accumulation using Droplet Digital PCR (ddPCR). The combination of these tools allows precise and reliable quantitation of resection intermediates^22^. We selected a locus on Chromosome I and analyzed ssDNA accumulation at 364bp from DSB ends in U-2OS cells^22,24^ (Fig. 2a). Importantly, all the ssDNA measurements through ddPCR were normalized on Cas9 cleavage efficiency to circumvent a potential decrease of cleavage efficiency in ATRi treated cells. Strikingly, VE-821 pretreatment caused a dose-dependent reduction in the ssDNA signal detected by ddPCR (Fig. 2b), consistent with the prediction that the depletion of resection proteins is causing loss of resection capacity. In particular, the highest VE-821 dose tested caused a decrease in ssDNA accumulation comparable to the profound loss of resection observed in cells where BRCA1 has been depleted by siRNA (Fig. 2c). To confirm that the observed impairment of resection required multi-day long-term ATRi treatment, and is not due to a rapid effect of the ATR inhibitor, such as impairment of protein-protein interactions^46^, we also measured ssDNA after an acute and high-dose VE-821 treatment. Consistent with the idea that the gradual depletion of central resection factors, and not a short-term effect of ATR inhibition, acute inhibition of ATR or CHK1 for 8 hours only caused a minor reduction in ssDNA exposure (Fig. 2d). Acute ATRi treatment did not cause any change in the abundance of BRCA1, CTIP, or BLM (Fig. 2e).

**Fig. 2.**
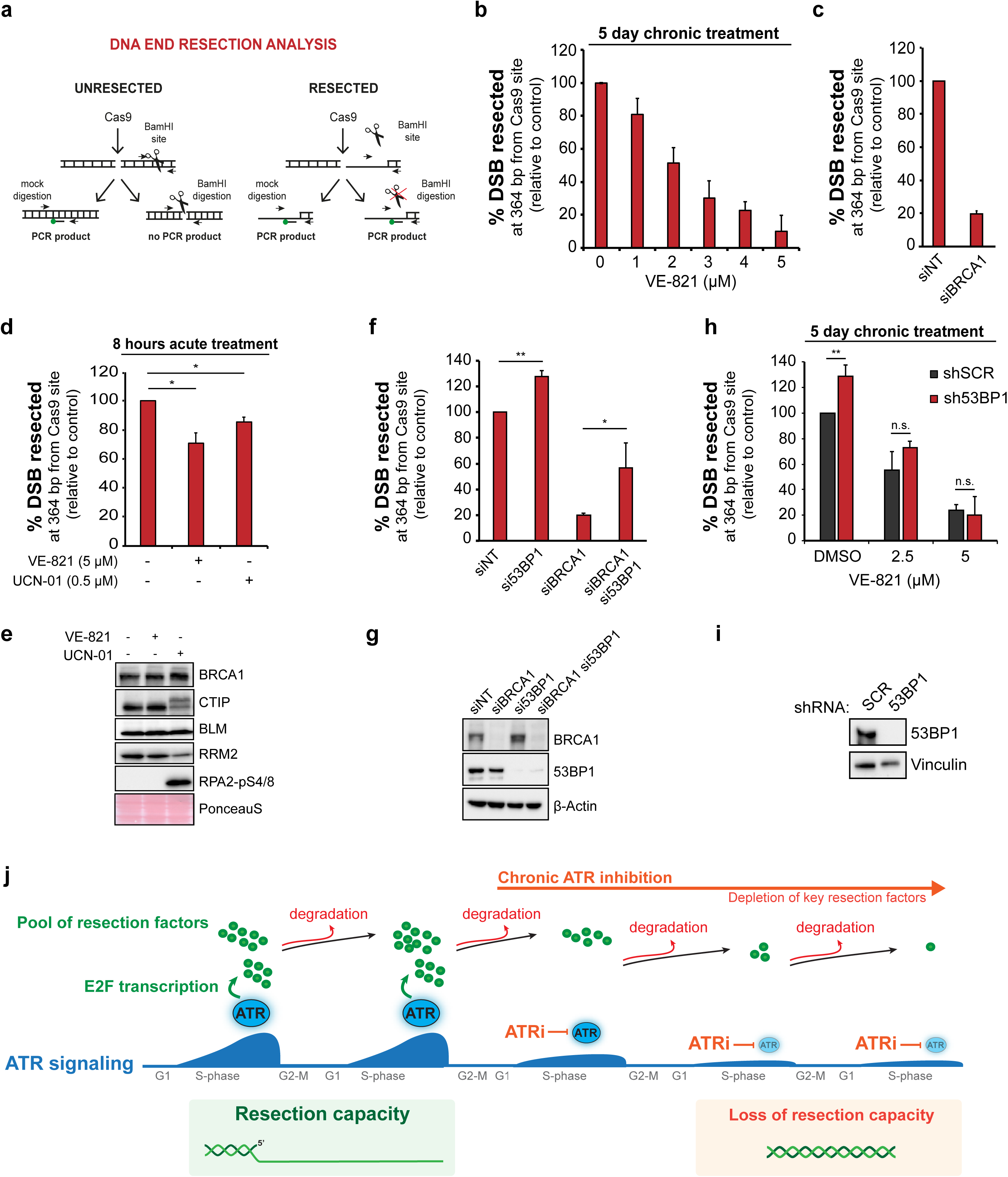
Long-term ATR inhibition severely impairs DNA end resection. (**a**) Experimental workflow of the CRISPR-Cas9-based resection assay used to induce DSBs in a defined locus on Chromosome I and the adopted restriction digestion strategy to measure ssDNA accumulation (**b**) U-2OS-SEC (Stably Expressing Cas9) cells were cultured for 5 days in medium containing DMSO or the indicated concentrations of VE-821 (ATRi). 24 hours prior to sgRNA transfection, Cas9-eGFP expression was induced by doxycycline (1*µ*g/ml). Cells were then harvested 8 hours after sgRNA transfection and processed for DNA extraction. Mean ± SD (n=4). (**c**) DNA end resection analysis in U-2OS-SEC 72 hours after transfection of siRNA against BRCA1. Results are same as shown in (**f**) (n=2) (**d**) DNA end resection analysis in U-2OS-SEC treated with 5*µ*M VE-821 (ATRi) or 0.5*µ*M UCN-01 (CHK1i) 8 hours after sgRNA transfection. Cas9-eGFP expression was induced 24 hours before sgRNA transfection. Mean ± SD (n=2). *P<0.05 (**e**) Immunoblot analysis of cells treated as in (**d**). (**f**) DNA end resection analysis in U-2OS-SEC 72 hours after transfection of the indicated siRNA. Mean ± SD (n=2). *P<0.05; **P<0.01 (**g**) Immunoblot analysis of cells treated as in (**f**). (**h**) DNA end resection analysis in U-2OS-SEC-shSCR and U-2OS-SEC-sh53BP1 cells treated for 5 days with the indicated VE-821 concentrations. After ATRi pretreatment, DSB was induced by co-transfecting sgRNA and purified Cas9. Mean ± SD (n=3). **P<0.01. (**i**) Immunoblot analysis of cells treated as in (**h**). (**j**) A schematic model showing how long-term ATRi treatment leads to the efficient depletion of HR proteins by preventing the *de novo* synthesis of new factors.

Because BRCA1 abundance is strongly affected by long-term ATR inhibition (Figs. 1a-d), we asked whether the impairment of resection was predominantly caused by the loss of BRCA1’s function in counteracting the anti-resection factor 53BP1. Since 53BP1 inactivation restores resection and HR in BRCA1-deficient tumors^47–49^, we asked whether loss of 53BP1 could restore resection in cells treated chronically with ATR inhibitors. Consistent with previous works, we found that 53BP1 depletion by siRNA significantly rescues resection in cells depleted for BRCA1, as measured by ddPCR at Cas9-induced breaks (Fig. 2f, g). Further analysis in U-2OS cells stably expressing inducible shRNA against 53BP1 and subjected to a 5-day pre-treatment with VE-821 revealed that 53BP1 inactivation does not accelerate resection speed upon long-term ATRi treatment (Fig. 2h, i). Therefore, loss of resection capacity in cells treated chronically with ATR inhibitors is not solely due to loss of BRCA1 but is likely a consequence of the loss of multiple important pro-resection factors. Overall, these results support the model whereby ATR inhibition severely impairs resection when cells undergo multiple cell divisions in the presence of ATR inhibitors (Fig. 2j). The long-term treatment not only prevents the *de novo* synthesis of resection proteins by blocking the E2F-mediated transcription of new factors^17^ but also allows for progressive degradation of the pre-existing pool (Fig. 2j). In addition, these findings establish a key pro-resection function for ATR, especially in cancer cells undergoing intrinsically high levels of ATR-CHK1 signaling due to elevated oncogene-induced replication stress. In these cells, increased ATR signaling should drive increased resection capacity and increased engagement of HR-mediated repair.

### Long-term ATR inhibition impairs RAD51 and RAD52 localization to DNA damage-induced foci

Given the differences in resection capacity upon long-term versus acute ATR inhibition, we reasoned that these distinct modes of ATRi treatment should lead to different outcomes in how DNA lesions are repaired. Acute ATR inhibition was previously reported to impair RAD51 localization to IR-induced foci, which requires ATR-mediated phosphorylation of PALB2 and its subsequent interaction with BRCA1^16^. Since resection is only mildly affected upon acute ATR inhibition, we predicted that cells treated acutely with ATR inhibitors should still be proficient in utilizing Single Strand Annealing (SSA), a homology-directed repair pathway dependent on RAD52, but independent of RAD51-PALB2-BRCA2^50^. Consistent with this prediction, we found that acute VE-821 treatment impaired PARPi-induced RAD51 foci in U-2OS cells (Fig. 3a, b), but did not alter PARPi-induced RAD52 foci (Fig. 3c, d). As a positive control for the requirement of resection for SSA^51^, the ability of cells to form PARPi-induced RAD52 foci was severely impaired in cells treated with the MRN inhibitor Mirin (Fig. 3d). Importantly, and congruent with the requirement of resection for SSA, long-term ATRi treatment severely reduced RAD52 foci formation after PARPi treatment (Fig. 3e, f). Long-term ATRi treatment further diminished the number of cells with detectable RAD51 foci compared to an acute ATRi treatment (Fig. 3g). These findings highlight how acute and long-term ATR treatment can drastically shape distinct DNA repair outcomes. In a condition where ATR signaling is not inhibited, ATR promotes HR by maintaining the proper abundance of the HR machinery and by directly phosphorylating HR factors (Fig. 3h). Since direct phosphorylation of HR factors by ATR seems dispensable for resection, but essential for RAD51 loading, acute ATR inhibition still allows resection, which in turn enables the engagement of resection-dependent repair pathways, such as RAD52-dependent repair (Fig. 3i). Long-term ATR inhibition leads to a distinct scenario, in which resection is severely blocked, therefore preventing both RAD51 and RAD52 engagement, and generally impairing any type of homology-directed repair (Fig. 3i).

**Fig. 3.**
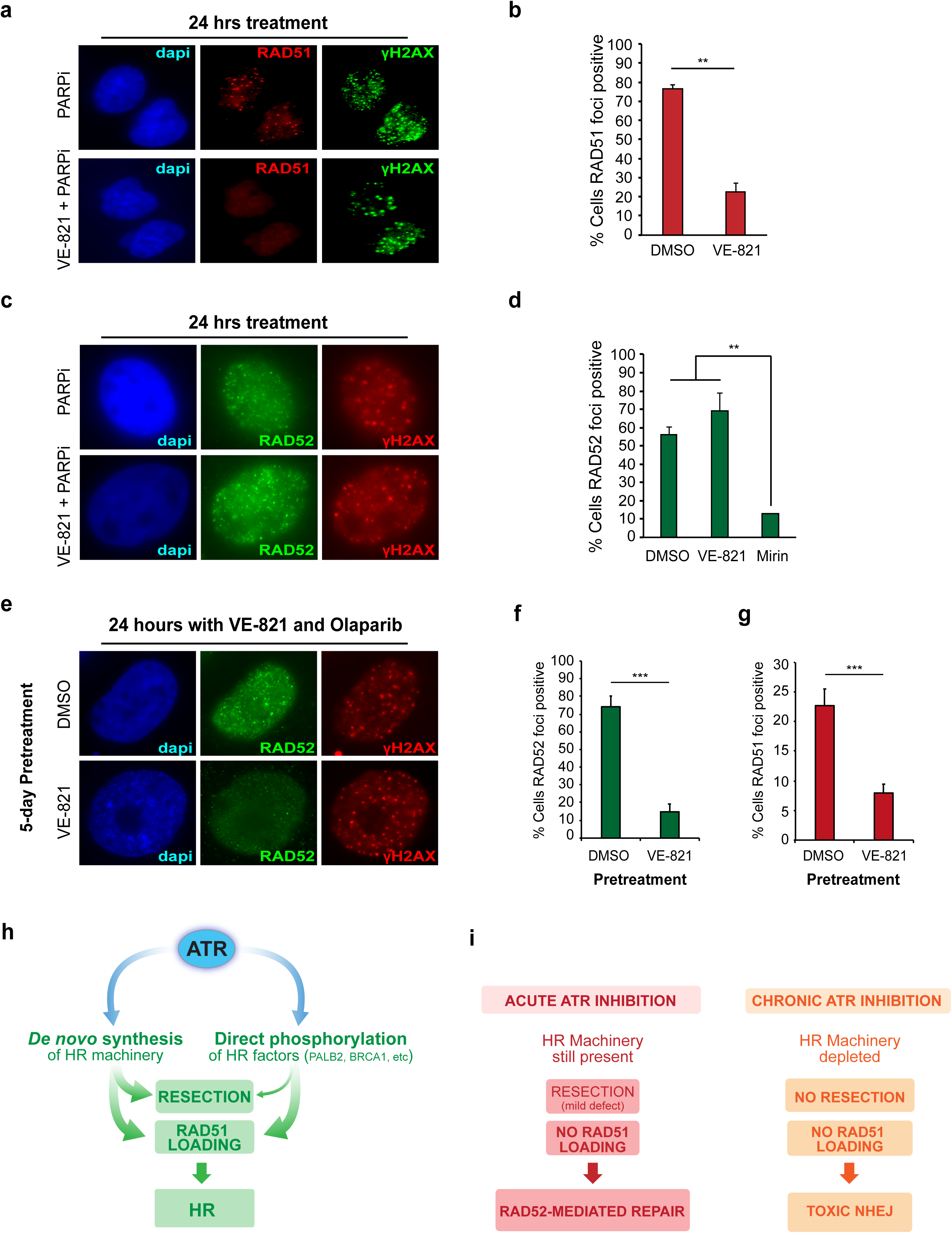
Long-term ATR inhibition impairs RAD51 and RAD52 localization. (**a**) Representative image showing RAD51 foci in U-2OS cells after a 24-hour treatment with Olaparib (10*µ*M) with or without VE-821 (2.5*µ*M) (**b**) Quantification of U-2OS cells as in (**a**) displaying >5 RAD51 distinct foci. Mean ± SD (n=3). **P<0.01 (**c**) Representative image showing RAD52 foci in U-2OS cells after a 24-hour treatment with Olaparib (10*µ*M) with or without VE-821 (2.5*µ*M) (**d**) Quantification of U-2OS cells as in (**c**) displaying RAD52 visible foci. Mean ± SD (n=3). **P<0.01. (**e**) Representative image showing RAD52 foci in U-2OS cells after a 5-day pretreatment with or without VE-821 (2.5*µ*M) and followed by a 24-hour treatment with Olaparib (10*µ*M) and VE-821 (2.5*µ*M). (**f**) Quantification of U-2OS cells treated as in (**e**) displaying RAD52 visible foci. Mean ± SD (n=3). ***P<0.001. (**g**) Quantification of U-2OS cells treated as in (**e**) displaying >5 RAD51 distinct foci. Mean ± SD (n=3). ***P<0.001. (**h**) ATR controls HR both through the *de novo* synthesis of DNA end resection and HR factors as well as through the direct phosphorylation of central HR proteins (*e.g.* BRCA1 and PALB2). (**i**) Distinct DNA repair outcomes upon acute or long-term ATR inhibition. While acute ATR inhibition impairs RAD51 loading but does not prevent DNA end resection allowing alternative RAD52-dependent DNA repair pathways, long-term ATR inhibition prevents both RAD51 and RAD52-dependent DNA repair enabling unscheduled NHEJ-repair.

### Long-term ATR inhibition induces hypersensitivity to PARP inhibitors in a DNA-PKcs-dependent manner

Since RAD51 or RAD52-dependent repair represent parallel HR pathways for promoting resistance to PARP inhibitors^50,52–55^, we reasoned that long-term ATR inhibition should lead to greater sensitization to PARP inhibitors as compared to acute ATR inhibition. To test this prediction, we monitored cell survival in cells subjected to a 5-day ATRi pre-treatment, followed by a 24-hour treatment with the PARP inhibitor Olaparib (Fig. 4a). At the end of the 5-day ATRi pre-treatment, cells should be highly defective in resection and unable to utilize any HR pathway for repairing PARPi-induced DNA lesions (Fig. 4a). We tested the protocol in three distinct cell lines, two cancer cell lines (HCT116 and U-2OS), and an untransformed cell line (RPE1). Congruent with our hypothesis, we found that ATRi pre-treatment increased the sensitivity of the two cancer cell lines to PARPi but did not increase sensitivity in the RPE1 cell line (Fig. 4b).

**Fig. 4.**
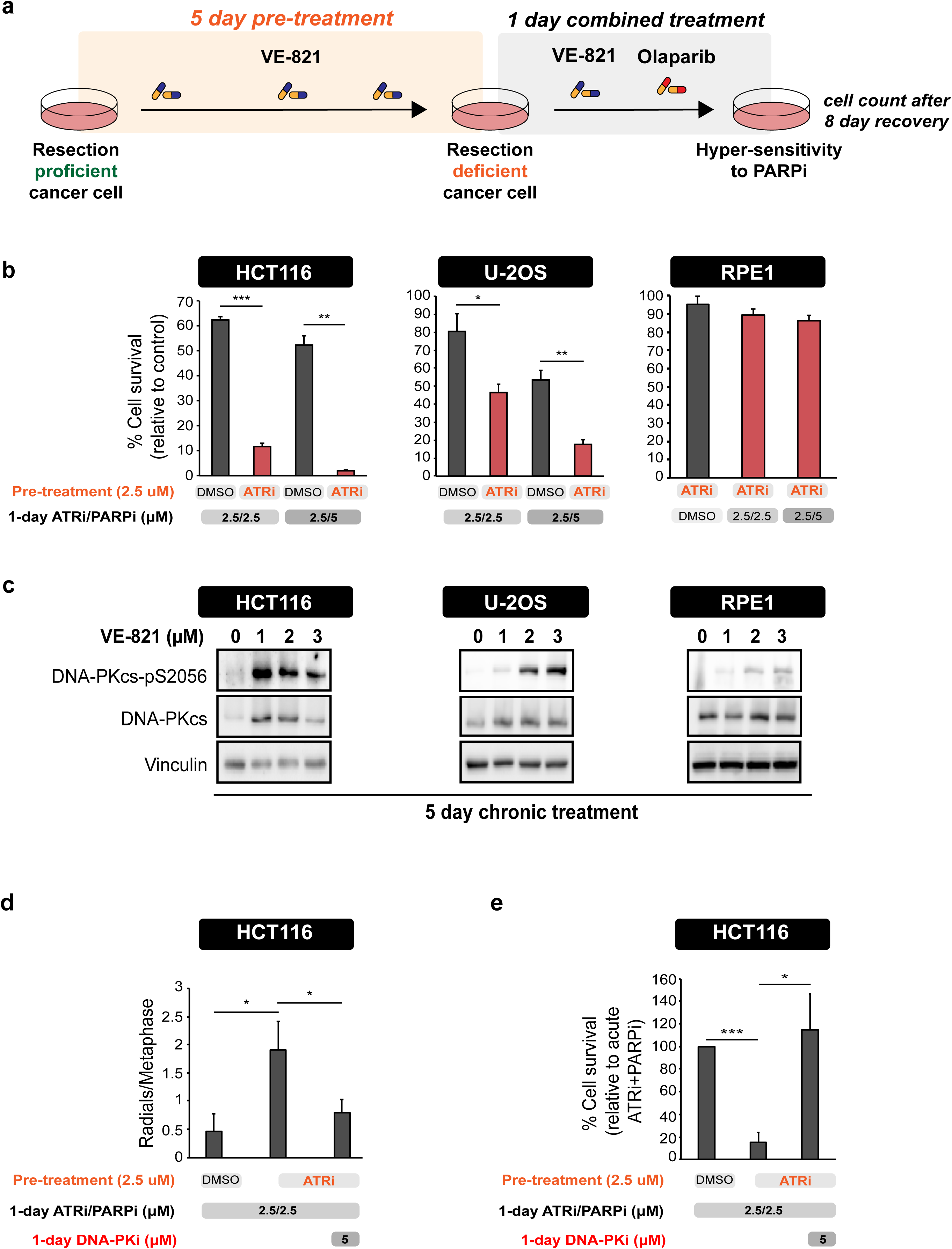
Long-term ATR inhibition induces hypersensitivity to PARPi and hyperactivation of DNA-PKcs in cancer cell lines. (**a**) Experimental workflow of the assay used to measure cell survival after long-term ATRi. (**b**) HCT116, U-2OS and RPE1 cells were treated as shown in the schematic (**a**). Cell viability was measured relative to cells treated with DMSO throughout all the pretreatment and treatment period. Mean ± SD (n=3). *P<0.05; **P<0.01; ***P<0.001. (**c**) Levels of DNA-PKcs-pS2056 and total DNA-PKcs in cells after a 5-day treatment with the indicated concentrations of VE-821. (**d**) HCT116 cells were treated as in (**a**) for 5 days. After 5 days, cells were treated with Olaparib (5*µ*M) with or without NU7441 (5*µ*M) for additional 24 hours. Metaphase spreads were then prepared as described in the Methods section. Mean ± SD (n=3). *P<0.05. (**e**) HCT116 cells were treated as in (**d**). Cell viability was measured relative to cells treated with acute ATRi and PARPi. Mean ± SD (n=4). *P<0.05; ***P<0.001.

Since loss or delayed DNA end resection leads to prolonged binding of NHEJ machinery at DSB ends^56^, we reasoned that long-term ATRi treatment leads to increased DNA-PKcs activation and, consequently, to the pronounced use of NHEJ to repair PARPi-induced lesions. Strikingly, long-term ATR inhibition induced activation of DNA-PKcs and the magnitude of DNA-PKcs activation in the different cell lines correlated with the degree of sensitization to PARPi conferred by the long-term ATRi pre-treatment (Fig. 4c).

The results suggest that the increased sensitization to PARPi induced by long-term ATRi pre-treatment is dependent on toxic NHEJ repair events that lead to loss of viability. Consistent with this possibility, we observed a significantly higher number of radial chromosomes, a typical output of toxic NHEJ of PARPi-induced breaks, in HCT116 cells exposed to the ATRi pre-treatment (Fig. 4d). Radial chromosomes after ATRi pre-treatment were NHEJ-dependent and where reduced by DNA-PK inhibition (Fig. 4d). In addition, the DNA-PKcs inhibitor NU7441 was able to rescue the sensitivity to PARPi conferred by the ATRi pre-treatment (Fig. 4e). Taken together, these findings support the model whereby ATRi pre-treatment induce hypersensitization to PARPi by allowing toxic NHEJ-mediated repair of DSBs. Of importance, the degree of sensitization to PARPi that is induced by long-term ATRi is distinct in different cell lines and directly correlates with the level of DNA-PKcs activation of each cell line.

### Long-term ATR inhibition bypasses overexpression of E2F1

E2F transcription is often elevated in cancer by mutations in the pRb pathway^57^. Since long-term ATR inhibition affects the abundance of E2F targets^17^, many of which are components of the HR machinery, the ability of cancer cells to upregulate E2F transcription could, in principle, bypass the effects of long-term ATR inhibition. In this case, cancer cells would become refractory to the effects of ATR inhibition and resection factor abundance would not decrease. To verify whether elevated E2F transcription could bypass the effects of long-term ATRi, we generated a conditional system to overexpress E2F1 using CRISPR/dCas9 transcriptional activation^58^. Using HCT116-dCas9-VP64 stable clones and five distinct sgRNAs targeting the E2F1 promoter, we were able to achieve robust overexpression of E2F1 (Fig. 5a, b). As expected, overexpression of E2F1 was associated with an increase in the abundance of several E2F targets (Fig. 5b). As a control, the abundance of 53BP1, whose expression is not regulated by E2F1, remained unaltered. Next, we subjected HCT116-dCas9-VP64 control or E2F1 overexpressing cells to a 5-day treatment with the ATRi AZD6738. Unexpectedly, long-term treatment with AZD6738 overcame high E2F1 expression and led to the efficient and gradual depletion of BRCA1 and CTIP resection factors (Fig. 5c). These results indicate that long-term ATRi treatment depletes DNA end resection factors independently of E2F1 status, suggesting that altering E2F expression does not represent a mechanism by which cancer cells may become refractory to changes in resection factors induced by long-term ATRi treatment.

**Fig. 5.**
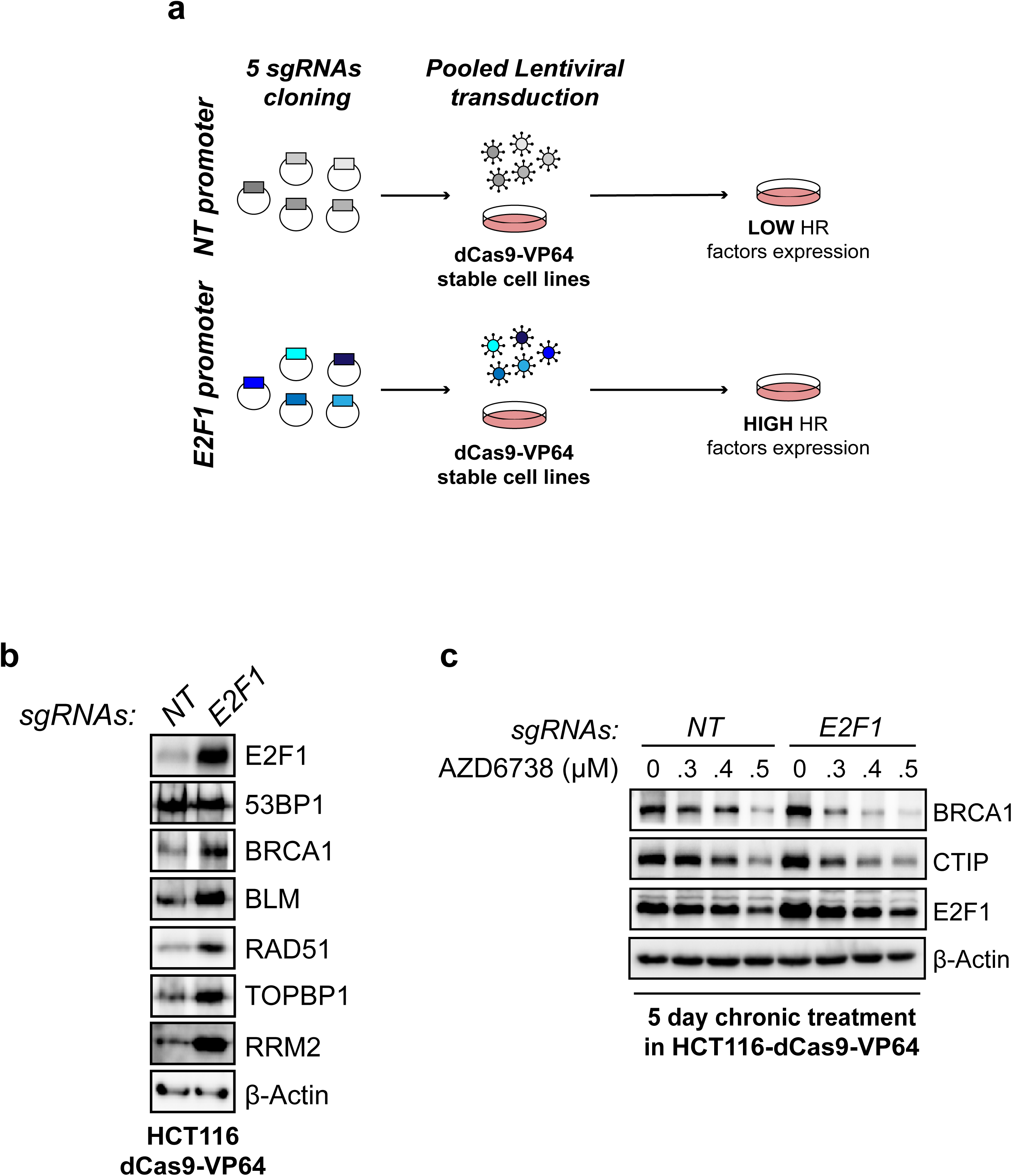
Long-term ATR inhibition bypasses overexpression of E2F1. (**a**) Schematic design of the CRISPR/dCas9 transcriptional activation protocol to induce E2F1 overexpression. (**b**) Immunoblot analysis of HCT116-dCas9-VP64 cells transduced with pooled lentivirus targeting a control region or the E2F1 promoter. (**c**) Immunoblot analysis of HCT116-dCas9-VP64 cells transduced with pooled lentivirus targeting or not the E2F1 promoter and treated for 5 days with the indicated concentrations of AZD6738.

## Discussion

5’-3’ DNA end resection is a crucial step in defining DNA repair outcomes. The ability to manipulate resection capacity is expected to be a powerful strategy to rationally modulate DNA repair outcomes in cancer cells and induce selective cell lethality. Here we report that ATR inhibitors can be used as potent modulators of DNA end resection. We further build on this finding to define how ATRi-induced resection loss promotes hypersensitivity to PARP inhibitors. Since PARP inhibitors are already FDA approved drugs used in cancer therapy, and ATR inhibitors are already in phase II clinical trials, our work should be directly applicable to the better design of drug treatment regimes. In particular, the understanding that long-term ATR inhibition promotes a resection block that forces NHEJ repair of PARPi-induced lesions reveals a defined rationale for enhancing the effectiveness of these inhibitors.

These findings establish a new role for ATR in modulating DNA end resection capacity. While previous studies elucidated the contribution of ATM and DNA-PKcs in resection initiation^24,59^, it was still unclear whether ATR contributes to DNA end resection. Part of this knowledge gap is caused by the employment of RPA or thymidine analog-based assays to visualize resection intermediates, which might conflict with the ATR role in suppressing origin firing and buffering RPA pool^60^. Here, employing a CRISPR-Cas9-based resection assay^22^, we found that up to 24 hours of ATR inhibition (acute inhibition) only caused a minor reduction in ssDNA accumulation. These findings are also in line with the established hierarchical mode of kinase activation at resected breaks, whereby ATR activation occurs after ATM signaling^26–30^. A modest reduction in ssDNA accumulation is still appreciable after acute ATR inhibition, indicating that ATR has a direct role in DNA end resection. Consistent with this idea, we have previously proposed that, upon recruitment and activation, ATR promotes an interaction between BRCA1 and the TOPBP1 scaffold to form a pro-resection complex that counteracts 53BP1-dependent resection inhibition^46^. While acute ATR inhibition has minor effects on resection, it does lead to severe reduction in RAD51 loading, as previous reported^16,61^, and confirmed here (Fig. 3a, b). Interestingly, however, the inability of acute ATRi treatment to impair DNA end resection allows the engagement of RAD52-dependent pathways that mitigate the cytotoxic effect of PARP inhibitors and other drugs. Previous studies have shown that RAD52-mediated repair could compensate for loss of HR in cells lacking the PALB2-BRCA2 machinery and therefore, could represent a mechanism of resistance to acute treatments with PARP inhibitors and ATR inhibitors^52,54,55,62^. Notably, RAD52 was reported to foster RAD51-dependent recombination in *brca2-*deficient cells^52^, therefore bypassing the requirement of PALB2 phosphorylation by ATR and possibly explaining the residual levels of RAD51 foci observable after acute ATRi and PARPi. Moreover, acute ATR inhibition has been shown to disrupt restored RAD51 foci and HR in *brca1 53bp1* cells after PARPi treatment^18^. Because *53bp1-* deficient cells have a pronounced preference for RAD52-mediated repair^63^, we predict that acute ATR inhibition would still allow RAD52-dependent pathways to occur in *brca1 53bp1* cells. In this sense, the ability of long-term ATRi treatments to inhibit both RAD51 and RAD52-dependent DNA repair provides the mechanistic rationale to utilize ATRi in treating HR-deficient tumors with acquired resistance to PARPi and other drugs. While it is still unclear whether the loss of a specific factor is responsible for the decrease in DNA end resection capacity upon long-term ATRi, our data suggest that loss of resection capacity upon long-term ATRi treatment is complex and due to loss of multiple pro-resection factors rather than loss of one specific factor, such as BRCA1. This is supported by our experiments showing that loss of 53BP1 does not increase resection efficiency upon long-term ATR inhibition. While loss of 53BP1 can restore HR and RAD51 foci in *brca1* cells^47^, it fails to do so in cells depleted of other resection factors such as CtIP^48,64^. Since the inability to promote resection upon long-term ATR inhibition could be due to the combined loss of BRCA1, CtIP, and additional pro-resection factors, we do not expect that restoring any specific factor alone is enough to restore resection.

Our findings strongly support the model whereby a decrease in DNA end resection and HR capacity upon long-term ATR inhibition leads to increased engagement of DNA-PKcs at DNA ends caused by PARPi, which promotes increased NHEJ, chromosomal aberrations, and cell death. Congruent with this model, we find that inhibition of DNA-PKcs restores cell viability in cells treated chronically with ATRi. This finding is also consistent with previous reports showing that genetic and chemical ablation of DNA-PKcs can suppress the formation of radial chromosomes in HR-deficient cancers and Fanconi Anemia patient-derived cells^65,66^. These reports and our findings raise the question of how cells deprived of both HR and NHEJ repair pathways survive to PARPi treatment. We propose that the engagement of DNA-PKcs at DNA breaks prevents any resection and rapidly directs DNA repair toward NHEJ, and that inhibition of DNA-PKcs allows time for residual resection to eventually occur, which in turn allows microhomology-mediated end joining (MMEJ) repair events. In the future, to understand the mechanism of DNA-PKcs-induced cell lethality, it would be essential to identify the DNA-PKcs target(s) through which DNA-PKcs promotes NHEJ. In particular, such analysis should be performed in the context of cells treated with ATRi, which might expand the spectrum of DNA-PKcs substrates, including proteins that are normally not targeted by the kinase. While other studies previously observed DNA-PKcs activation after ATR inhibition^12,67^, it is still unclear what DNA structures arising after ATR inhibition trigger DNA-PKcs activation in cells. A possible scenario is that the gradual decrease in resection capacity caused by long-term suppression of ATR signaling leads to a progressive engagement of DNA-PKcs at DNA breaks naturally forming during DNA replication or as a consequence of ATR inhibition. Also, DNA breaks could arise from extensive degradation of stalled replication forks, since long-term ATR inhibition could deplete the abundance of fork protection factors such as BRCA1/2^17^. As a consequence, extensive fork degradation allows the SLX4-MUS81 nuclease complex to cleave replication forks leading to DNA-PKcs activation^12,68,69^. According to our data, DNA-PKcs activation by long-term ATRi pretreatment is not only a key event in the sensitization of cancer cells to PARP inhibitors but also the magnitude of DNA-PKcs activation/signaling could be considered as a predictive marker of long-term ATRi efficacy in sensitizing cancer cells to PARP inhibitors. These findings provide mechanistic rationales for the design of more effective inhibitor treatment regimes and to better predict the treatment efficiency based on the levels of DNA-PKcs activation.

Modulation of the E2F-dependent transcription program is at the heart of the observed effects of long-term ATR inhibition. E2F-dependent transcription has a range of relevant implications to understand cancer proliferation and the ability of cancer cells to withstand high levels of endogenous and genotoxin-induced replication stress. E2F transcription is increased by oncogenic mutations that alleviate the pRb inhibitory activity on E2F1 (mutations on pRb or CDKN2A, for instance). In addition, increased E2F-transcription is a common feature of cancers undergoing increased levels of intrinsic replication stress and ATR signaling^32,70^. We further speculate that E2F transcription is also induced in tumors undergoing repeated cycles of chemotherapy, and that increases in the E2F-dependent transcriptional program represent a potential source of resistance to anticancer drugs. In agreement with this idea, a recent study showed that a pre-exposure to cisplatin is sufficient to induce an ATR-dependent adaptive response to subsequent cisplatin treatments that involves transcription of the PRIMPOL protein to rescue fork degradation in *brca1* cancer cells^71^. This finding is in line with our model positioning ATR as a sensor of intrinsic or drug-induced replication stress and a regulator of genome stability through the modulation of the E2F transcription program and HR-coupled repair. Importantly, our ability to overcome induced E2F1 overexpression by long-term ATR inhibition indicates that resection and HR capacity can be remodeled independently of the levels of E2F transcription. This finding should be relevant to addressing ATRi-mediated therapies in tumors that are highly dependent on E2F transcription or in tumors that have built an adaptive response to chemotherapy^71,72^.

In the future, it will be essential to transfer the knowledge accumulated about the effect of long-term ATR inhibition to more complex systems such as tumor organoids and, more importantly, to mouse models of human cancers. To date, an effective strategy to inhibit DNA end resection with high tolerability and minimized side effects in patients is still missing. Long-term ATR inhibition represents an innovative and efficient strategy to inhibit DNA end resection and manipulate DNA repair outcomes in many cancers.

## Methods

### Cells

U-2OS, HCT116, RPE1, 293T cells were cultured in Dulbecco Modified Medium supplemented with 10% fetal calf serum, 1% penicillin/streptomycin and 1% Non-essential aminoacids. HCT116-*ETAA1ΔAAD*-TOPBP1-mAID, a kind gift from David Cortez, were cultured in Dulbecco Modified Medium supplemented with 10% fetal calf serum, 1% penicillin/streptomycin and 1% Non-essential aminoacids. U-2OS-SEC (Stably Expressing inducible Cas9) clones were generated by lentiviral infection with TLCV2 vector (a kind gift from Adam Karpf, Addgene plasmid #87360) followed by puromycin selection (1*µ*g/ml). HCT116-dCas9-VP64 clones were generated by lentiviral infection with the pHAGE EF1α dCas9-VP64 vector (a kind gift from Rene Maehr and Scot Wolfe, Addgene plasmid #50918) followed by puromycin selection (1*µ*g/ml).

U-2OS-shSCRAMBLE and U-2OS-sh53BP1 were generated by lentiviral infection with pLKO.1 derivative plasmid followed by puromycin selection (1*µ*g/ml). shSCRAMBLE.FOR: CCGGCCTAAGGTTAAGTCGCCCTCGCTCGAGCGAGGGCGACTTAACCTTAGGTTTTTG.

shSCRAMBLE.REV:AATTCAAAAACCTAAGGTTAAGTCGCCCTCGCTCGAGCGAGGGCGACTTAA CCTTAGG.

sh53BP1.FOR:CCGGGATACTCCTTGCCTGATAATTCTCGAGAATTATCAGGCAAGGAGTATCTTTTTG.

sh53BP1.REV:ATTCAAAAAGATACTCCTTGCCTGATAATTCTCGAGAATTATCAGGCAAGGAGTATC.

All the cell lines were regularly tested for mycoplasma contamination with the Universal Mycoplasma Detection Kit (ATCC).

### Inhibitors and chemicals

The inhibitors used in this study are VE-821 (ATRi, Selleckchem), AZD6738 (ATRi, Selleckchem), UCN-01 (CHK1i, Sigma Millipore), Mirin (MRE11i, Sigma Millipore), NU7441 (DNA-PKi, Selleckchem), Olaparib (PARPi, Selleckchem). 5-iododeoxyuridine (IdU, Sigma Millipore) was used at 25μM concentration. Auxin (IAA, Sigma Millipore) was used at a 10*µ*g/ml concentration.

### Antibodies

The antibodies used in this study are: BRCA1^73^ (provided by Raimundo Freire), CTIP (A300-488A, Bethyl Laboratories), BLM (A300-110A, Bethyl Laboratories), 53BP1 (NB100-304, Novus Biologicals), β-Actin (MA1-140, Thermo Fisher Scientific), TOPBP1^73^ (provided by Raimundo Freire), RRM2^17^ (provided by Raimundo Freire), RPA2-pS4/8 (A300-245A, Bethyl Laboratories), DNA-PKcs-pS2056 (PA5-78130, Thermo Fisher Scientific), DNA-PKcs (A300-516A-T, Bethyl Laboratories), Vinculin (#4650, Cell signaling), RAD51 (PC130, Calbiochem), RAD52 (5E11E7, Thermo Fisher Scientific), γH2AX (JBW301, Sigma Millipore), γH2AX (A300-081A, Bethyl Laboratories), E2F1 (sc-251, Santa Cruz Biotechnology).

### RNAi

U-2OS-SEC cells were transfected with the indicated siRNA using Lipofectamine RNAiMAX (Thermo Fisher Scientific) according to the manufacturer’s instructions and used 72 hours after siRNA transfection. siNT was purchased from Ambion (Cat#AM4629), siBRCA1: AAAUGUCACUCUGAGAGGAUAGCCC, si53BP1: AGAACGAGGAGACGGUAAUAGUGGG.

### Cell Cycle Analysis

To analyze cell cycle distribution, cells were pulse-labelled with 25*µ*M IdU for 30 minutes. After fixation, an additional incubation with BrdU primary antibody followed by an incubation with AlexaFluor488 secondary antibody was done. Data acquisition was performed with a FACS DIVA Software.

### DSB generation through CRISPR-Cas9

For resection experiments in U-2OS-SEC, DSB2 sgRNAs were synthetized and purchased from Thermo Fisher Scientific and transfected using Lipofectamine RNAiMAX (Thermo Fisher Scientific) according to the manufacturer’s instructions. Prior to sgRNA transfection, Cas9-eGFP expression was induced for 24 hours with 1*µ*g/ml doxycycline. For resection experiments in Cas9 not expressing cell lines, DSB2 sgRNAs and TrueCut Cas9 protein were synthetized and purchased from Thermo Fisher Scientific and transfected using Lipofectamine CRISPRMAX (Thermo Fisher Scientific) according to the manufacturer’s instructions.

### Genomic DNA extraction

U-2OS cells were pretreated with ATRi VE-821 and then seeded O/N on 12-well plate. 8 hours after sgRNA transfection, cells were harvested and genomic DNA was extracted by Nucleospin™ Tissue Kit (Macherey-Nagel) according to the manufacturer’s instructions. The day after, a desired volume of genomic DNA was equally mock or digested with *Bam*HI (New England BioLabs) for 4 h at 37°C. Digested and mock digested DNA was precipitated, purified and 5*µ*l were used for each ddPCR reaction.

### DNA end resection measurement through Droplet Digital PCR (ddPCR)

The ddPCR reaction was assembled as follows: 5*µ*l of genomic DNA, 1X ddPCR™ Supermix for Probes (no dUTP, Bio-Rad), 900nM for each pair of primers, 250nM for each probe, and dH_2_O to 20*µ*l per sample. Droplets were produced pipetting 20*µ*l of the PCR reaction mix into single wells of a universal DG8™ cartridge^®^ for droplets generation (Bio-Rad). 70*µ*l of droplet generation oil^®^ was also added in each well next to the ones containing the samples. Cartridges were covered with DG8™ droplet generator gaskets (Bio-Rad) and then placed into the droplet generator (QX200™, Bio-Rad). After droplet generation, 40*µ*l of emulsion were transferred from the cartridge to a 96-well ddPCR plate (Bio-Rad). Before PCR reaction, 96-well PCR plates were sealed with peelable foil heat seals at the PCR plate sealer machine (PX1™, Bio-Rad). For PCR reaction, Taq polymerase was activated at 95°C for 5 minutes and then 39 cycles of 95°C for 30 s and 58.7°C for 1 minute were made. At the end of the cycles, a 5 minute-step at 90°C was made and then temperature was held at 12°C. After the PCR, FAM and HEX fluorescence was read at the droplet reader (QX200™, Bio-Rad) using QuantaSoft™ software (Bio-Rad). For each sample the number of droplets generated were on an average of 15,000. The number of copies/*µ*l of the target loci was determined setting an empirical baseline threshold identical in all the samples. For the calculation of Cas9 cleavage efficiency, a ratio (*r*) was made between the number of copies of the locus across the Cas9 site (HEX probe) and a control locus on Chr. XXII (FAM probe) in cells transfected or not with the sgRNA. We then calculated R= *r*_+gRNA_/*r*_-gRNA_ and the final cleavage efficiency with the following equation: % Cas9 cleavage efficiency= (1-R)*100. For the measurement of ssDNA generated by the resection process (% ssDNA) we calculated the ratio (*r’*) between the number of copies of DSB2 locus (364bp from the Cas9 site) and a control locus on Chr. XXII with or without sgRNA digested or mock with *Bam*HI restriction enzyme. The absolute percentage of ssDNA was then calculated with the following equation: % ssDNA= (*r’*_digested_/*r’*_mock_)_+gRNA_-(*r’*_digested_/*r’*_mock_)_-gRNA_. The final percentage of DSB resected was calculated making the ratio between the % ssDNA and the % Cas9 cleavage efficiency.

### E2F1 overexpression through CRISPRa

Five different sgRNAs sequences targeting the E2F1 promoter were individually cloned into a LentisgRNA-neo (a gift from Brett Stringer, Addgene plasmid #104992). sgRNAs were designed using the GPP sgRNA designer tool, from Broad Institute/MIT (https://portals.broadinstitute.org/gpp/public/analysis-tools/sgrna-design-crisprai) (sequences are available in the Supplementary Table 1). Lentivirus for each sgRNA were produced in 293T cells using standard procedures. HCT116-dCas9-VP64 stable clone was transduced with viral pools containing five different sgRNAs specific for the E2F1 promoter. Viral transduction was then followed by selection with G418 (700*µ*g/ml) for 3-5 days.

### Cell viability assay

U-2OS, HCT116, or RPE1 cells were treated for 5 days with DMSO or 2.5 *µ*M VE-821, refreshing media on days 2 and 4. Then either 1 × 10^5^ cells were passaged in new plates with media containing DMSO, VE-821 (2.5 *µ*M), Olaparib (2.5 *µ*M or 5 *µ*M), or a combination of VE-821 and Olaparib. Cells were treated in these conditions for 24 hours, after which the drugged media was removed, allowing the cells to recover for 8 days in drug-free media. After the recovery period, live cell number was quantified. Live cell number quantification was performed by trypsinizing the cells and counting with the MOXI Z Automated Cell Counter Kit (Orflo, MXZ001).

### Metaphase spreads preparation

Prior to harvest, cells were treated with 150 ng/ml Colcemid for 1 hour and then collected by centrifugation. Cell pellets were shortly resuspended in Hypotonic Buffer and then fixed in fixation buffer overnight (3:1 Methanol: Acetic Acid). Fixed cells were extensively washed with fixation buffer and then spotted on microscope slides with Vectashield Antifade mounting medium with DAPI (Vector Laboratories). Metaphase spreads were imaged using a Leica DFC9000 GTC cMOS camera with a 100X objective. Each condition was repeated in three independent biological experiments approximately 50 metaphases were analyzed per condition. The two-tailed Student’s t test was used for statistical analysis.

### Immunofluorescence and Microscopy Analysis

U-2OS cells were grown on coverslips and treated with the indicated combination of acute/chronic ATRi and PARPi treatment. Cells were then fixed with 3.7% formaldehyde in PBS for 10min at RT. Fixed cells were then washed three times with PBS, permeabilized 5 minutes with 0,2% Triton-X100/PBS at RT and blocked in 10%BSA/PBS 20 minutes at RT. Coverslips were incubated first with primary antibodies for 2 hours at RT, followed by three washes with PBS, and then for 1 hour with relative secondary antibodies (Alexa Fluor488 Goat anti-Mouse IgG (H+L) and Alexa Fluor568 Donkey anti-Rabbit IgG (H+L), Thermo Fisher Scientific). After incubation with secondary antibody, coverslips were washed three times with PBS and then mounted on glass microscope slides using DAPI-Vectashield mounting medium (Vector Laboratories). Microscope slides were imaged using a Leica DMi8 inverted fluorescent microscope with a 63X objective. For RAD51 and RAD52 foci scoring, approximately 150-200 cells/replicate were counted and the fraction of cells with more than 5 distinct RAD51 foci or 10 distinct RAD52 foci was determined. The two-tailed Student’s t test was used for statistical analysis.

### Immunoblotting analysis

Cells were harvested and lysed in modified RIPA buffer (50mM Tris-HCl pH 7.5, 150mM NaCl, 1% Tergitol, 0.25% Sodium Deoxycholate, 5mM EDTA) supplemented with Complete EDTA-free protease inhibitor cocktail (Roche), 1mM PMSF and 5mM NaF. Whole cell lysates, after sonication, were cleared by 15 minutes centrifugation at 13,000 rpm at 4°C. 20*µ*g of protein extract were mixed with 3X SDS Sample Buffer and resolved by SDS-PAGE. Gel were transferred on PVDF membranes and western blot signal was acquired with a Chemidoc Imaging System (Bio-Rad).

### Statistical analysis

All experimental results were analyzed using unpaired two-tailed Student’s t test as indicated in figure legends.

## Data availability

The authors declare that all data supporting the findings of this study are available within the article or from the corresponding author upon request.

## Author contribution

D.D. and M.B.S. designed the study. D.D., J.R.S, C.F.R.A and K.F. performed the experiments and analyzed data. R.F. provided critical reagents. D.D. and M.B.S wrote the manuscript.

## Acknowledgements

We thank David Cortez for useful reagents. We thank Fenghua Hu and Tony Bretscher for the use of the Microscopes. We thank Beatriz Almeida for technical assistance. We thank all the members of the Smolka lab for helpful discussion. D.D. is supported by a fellowship from the Fleming Research Foundation. This work was supported by M.B.S. grants from the National Institutes of Health (R01GM097272).

## REFERENCES

1. Cortez, D. Replication-Coupled DNA Repair. Mol. Cell 74, 866–876 (2019).

2. Cimprich, K. A. & Cortez, D. ATR: An essential regulator of genome integrity. Nat. Rev. Mol. Cell Biol. 9, 616–627 (2008).

3. Zou, L. & Elledge, S. J. Sensing DNA damage through ATRIP recognition of RPA-ssDNA complexes. Science (80-.). 300, 1542–1548 (2003).

4. Bass, T. E. et al. ETAA1 acts at stalled replication forks to maintain genome integrity. Nat. Cell Biol. 18, 1185–1195 (2016).

5. Haahr, P. et al. Activation of the ATR kinase by the RPA-binding protein ETAA1. Nat. Cell Biol. 18, 1196–1207 (2016).

6. Kumagai, A., Lee, J., Yoo, H. Y. & Dunphy, W. G. TopBP1 activates the ATR-ATRIP complex. Cell 124, 943–955 (2006).

7. Mordes, D. A., Glick, G. G., Zhao, R. & Cortez, D. TopBP1 activates ATR through ATRIP and a PIKK regulatory domain. Genes Dev. 22, 1478–1489 (2008).

8. Lee, Y. C., Zhou, Q., Chen, J. & Yuan, J. RPA-Binding Protein ETAA1 Is an ATR Activator Involved in DNA Replication Stress Response. Curr. Biol. 26, 3257–3268 (2016).

9. Guo, Z., Kumagai, A., Wang, S. X. & Dunphy, W. G. Requirement for Atr in phosphorylation of Chk1 and cell cycle regulation in response to DNA replication blocks and UV-damaged DNA in Xenopus egg extracts. Genes Dev. 14, 2745–2756 (2000).

10. Hui, Z. & Helen, P.-W. ATR-Mediated Checkpoint Pathways Regulate Phosphorylation and Activation of Human Chk1. Mol. Cell. Biol. 21, 4129–4139 (2001).

11. Liu, Q. et al. Chk1 is an essential kinase that is regulated by Atr and required for the G2/M DNA damage checkpoint. Genes Dev. 14, 1448–1459 (2000).

12. Buisson, R., Boisvert, J. L., Benes, C. H. & Zou, L. Distinct but Concerted Roles of ATR, DNA-PK, and Chk1 in Countering Replication Stress during S Phase. Mol. Cell 59, 1011–1024 (2015).

13. Toledo, L. I. et al. A cell-based screen identifies ATR inhibitors with synthetic lethal properties for cancer-associated mutations. Nat. Struct. Mol. Biol. 18, 721–727 (2011).

14. Murga, M. et al. Exploiting oncogene-induced replicative stress for the selective killing of Myc-driven tumors. Nat. Struct. Mol. Biol. 18, 1331–1335 (2011).

15. Brown, E. J. & Baltimore, D. Essential and dispensable roles of ATR in cell cycle arrest and genome maintenance. Genes Dev. 17, 615–628 (2003).

16. Buisson, R. et al. Coupling of Homologous Recombination and the Checkpoint by ATR. Mol. Cell 65, 336–346 (2017).

17. Kim, D., Liu, Y., Oberly, S., Freire, R. & Smolka, M. B. ATR-mediated proteome remodeling is a major determinant of homologous recombination capacity in cancer cells. Nucleic Acids Res. 46, 8311–8325 (2018).

18. Yazinski, S. A. et al. ATR inhibition disrupts rewired homologous recombination and fork protection pathways in PARP inhibitor-resistant BRCA-deficient cancer cells. Genes Dev. 31, 318–332 (2017).

19. Symington, L. S. Mechanism and regulation of DNA end resection in eukaryotes. Crit. Rev. Biochem. Mol. Biol. 51, 195–212 (2016).

20. Chen, L., Nievera, C. J., Lee, A. Y. L. & Wu, X. Cell cycle-dependent complex formation of BRCA1·CtIP·MRN is important for DNA double-strand break repair. J. Biol. Chem. 283, 7713–7720 (2008).

21. Cruz-García, A., López-Saavedra, A. & Huertas, P. BRCA1 accelerates CtIP-ediated DNA-end resection. Cell Rep. 9, 451–459 (2014).

22. Dibitetto, D., La Monica, M., Ferrari, M., Marini, F. & Pellicioli, A. Formation and nucleolytic processing of Cas9-induced DNA breaks in human cells quantified by droplet digital PCR. DNA Repair (Amst). 68, (2018).

23. Sartori, A. A. et al. Human CtIP promotes DNA end resection. Nature 450, 509–514 (2007).

24. Zhou, Y., Caron, P., Legube, G. & Paull, T. T. Quantitation of DNA double-strand break resection intermediates in human cells. Nucleic Acids Res. 42, 1–11 (2014).

25. Gravel, S., Chapman, J. R., Magill, C. & Jackson, S. P. DNA helicases Sgs1 and BLM promote DNA double-strand break resection. Genes Dev. 22, 2767–2772 (2008).

26. Adams, K. E., Medhurst, A. L., Dart, D. A. & Lakin, N. D. Recruitment of ATR to sites of ionising radiation-induced DNA damage requires ATM and components of the MRN protein complex. Oncogene 25, 3894–3904 (2006).

27. Cuadrado, M. et al. ATM regulates ATR chromatin loading in response to DNA double-strand breaks. J. Exp. Med. 203, 297–303 (2006).

28. Jazayeri, A. et al. ATM- and cell cycle-dependent regulation of ATR in response to DNA double-strand breaks. Nat. Cell Biol. 8, 37–45 (2006).

29. Myers, J. S. & Cortez, D. Rapid activation of ATR by ionizing radiation requires ATM and Mre11. J. Biol. Chem. 281, 9346–9350 (2006).

30. Shiotani, B. & Zou, L. Single-Stranded DNA Orchestrates an ATM-to-ATR Switch at DNA Breaks. Mol. Cell 33, 547–558 (2009).

31. Buisson, R. et al. Cooperation of breast cancer proteins PALB2 and piccolo BRCA2 in stimulating homologous recombination. Nat. Struct. Mol. Biol. 17, 1247–1254 (2010).

32. Bertoli, C., Klier, S., McGowan, C., Wittenberg, C. & De Bruin, R. A. M. Chk1 inhibits E2F6 repressor function in response to replication stress to maintain cell-cycle transcription. Curr. Biol. 23, 1629–1637 (2013).

33. Yuan, R. et al. Chk1 and 14-3-3 proteins inhibit atypical E2Fs to prevent a permanent cell cycle arrest. The EMBO Journal vol. 37 (2018).

34. Charrier, J. D. et al. Discovery of Potent and Selective Inhibitors of Ataxia Telangiectasia Mutated and Rad3 Related (ATR) Protein Kinase as Potential Anticancer Agents. J. Med. Chem. 54, 2320–2330 (2011).

35. Karnitz, L. M. & Zou, L. Molecular pathways: Targeting ATR in cancer therapy. Clin. Cancer Res. 21, 4780–4785 (2015).

36. Lecona, E. & Fernandez-Capetillo, O. Targeting ATR in cancer. Nat. Rev. Cancer 18, 586–595 (2018).

37. Rundle, S., Bradbury, A., Drew, Y. & Curtin, N. J. Targeting the ATR-CHK1 axis in cancer therapy. Cancers (Basel). 9, 1–25 (2017).

38. Mei, L., Zhang, J., He, K. & Zhang, J. Ataxia telangiectasia and Rad3-related inhibitors and cancer therapy: Where we stand. J. Hematol. Oncol. 12, 1–8 (2019).

39. Foote, K. M. et al. Discovery and Characterization of AZD6738, a Potent Inhibitor of Ataxia Telangiectasia Mutated and Rad3 Related (ATR) Kinase with Application as an Anticancer Agent. J. Med. Chem. 61, 9889–9907 (2018).

40. Kim, H. et al. Targeting the ATR/CHK1 axis with PARP inhibition results in tumor regression in BRCA-mutant ovarian cancer models. Clin. Cancer Res. 23, 3097–3108 (2017).

41. Ning, J. F. et al. Myc targeted CDK18 promotes ATR and homologous recombination to mediate PARP inhibitor resistance in glioblastoma. Nat. Commun. 10, 2910 (2019).

42. Schoonen, P. M. et al. Premature mitotic entry induced by ATR inhibition potentiates olaparib inhibition-mediated genomic instability, inflammatory signaling, and cytotoxicity in BRCA2-deficient cancer cells. Mol. Oncol. 13, 2422–2440 (2019).

43. Wengner, A. M. et al. The novel ATR inhibitor BAY 1895344 is efficacious as monotherapy and combined with DNA damage-inducing or repair-compromising therapies in preclinical cancer models. Mol. Cancer Ther. molcanther.0019.2019 (2019) doi:10.1158/1535-7163.mct-19-0019.

44. Bass, T. E. & Cortez, D. Quantitative phosphoproteomics reveals mitotic function of the ATR activator ETAA1. J. Cell Biol. 218, 1235–1249 (2019).

45. Saldivar, J. C. et al. An intrinsic S/G2 checkpoint enforced by ATR. Science (80-.). 361, 806–810 (2018).

46. Liu, Y. et al. TOPBP1Dpb11 plays a conserved role in homologous recombination DNA repair through the coordinated recruitment of 53BP1Rad9. J. Cell Biol. 216, (2017).

47. Bouwman, P. et al. 53BP1 loss rescues BRCA1 deficiency and is associated with triple-negative and BRCA-mutated breast cancers. Nat. Struct. Mol. Biol. 17, 688–695 (2010).

48. Bunting, S. F. et al. 53BP1 inhibits homologous recombination in brca1-deficient cells by blocking resection of DNA breaks. Cell 141, 243–254 (2010).

49. Callen, E. et al. 53BP1 Enforces Distinct Pre- and Post-resection Blocks on Homologous Recombination. Mol. Cell 1–13 (2019) doi:10.1016/j.molcel.2019.09.024.

50. Stark, J. M., Pierce, A. J., Oh, J., Pastink, A. & Jasin, M. Genetic Steps of Mammalian Homologous Repair with Distinct Mutagenic Consequences. Mol. Cell. Biol. 24, 9305–9316 (2004).

51. Bhargava, R., Onyango, D. O. & Stark, J. M. Regulation of Single-Strand Annealing and its Role in Genome Maintenance. Trends Genet. 32, 566–575 (2016).

52. Feng, Z. et al. Rad52 inactivation is synthetically lethal with BRCA2 deficiency. Proc. Natl. Acad. Sci. U. S. A. 108, 686–691 (2011).

53. Wray, J., Liu, J., Nickoloff, J. A. & Shen, Z. Distinct RAD51 associations with RAD52 and BCCIP in response to DNA damage and replication stress. Cancer Res. 68, 2699–2707 (2008).

54. Chandramouly, G. et al. Small-Molecule Disruption of RAD52 Rings as a Mechanism for Precision Medicine in BRCA-Deficient Cancers. Chemistry and Biology vol. 22 1491–1504 (2015).

55. Sullivan-Reed, K. et al. Simultaneous Targeting of PARP1 and RAD52 Triggers Dual Synthetic Lethality in BRCA-Deficient Tumor Cells. Cell Rep. 23, 3127–3136 (2018).

56. Langerak, P., Mejia-Ramirez, E., Limbo, O. & Russell, P. Release of Ku and MRN from DNA ends by Mre11 nuclease activity and Ctp1 is required for homologous recombination repair of double-strand breaks. PLoS Genet. 7, (2011).

57. Nevins, J. R. The Rb/E2F pathway and cancer. Hum. Mol. Genet. 10, 699–703 (2001).

58. Konermann, S. et al. Genome-scale transcriptional activation by an engineered CRISPR-Cas9 complex. Nature 517, 583–588 (2015).

59. Zhou, Y. & Paull, T. T. DNA-dependent protein kinase regulates DNA end resection in concert with Mre11-Rad50-Nbs1 (MRN) and Ataxia Telangiectasia-mutated (ATM). J. Biol. Chem. 288, 37112–37125 (2013).

60. Toledo, L. I. et al. ATR prohibits replication catastrophe by preventing global exhaustion of RPA. Cell 155, 1088 (2013).

61. Sørensen, C. S. et al. The cell-cycle checkpoint kinase Chk1 is required for mammalian homologous recombination repair. Nat. Cell Biol. 7, 195–201 (2005).

62. Lok, B. H., Carley, A. C., Tchang, B. & Powell, S. N. RAD52 inactivation is synthetically lethal with deficiencies in BRCA1 and PALB2 in addition to BRCA2 through RAD51-mediated homologous recombination. Oncogene 32, 3552–3558 (2013).

63. Ochs, F. et al. 53BP1 fosters fidelity of homology-directed DNA repair. Nat. Struct. Mol. Biol. 23, 714–721 (2016).

64. Polato, F. et al. CtIP-mediated resection is essential for viability and can operate independently of BRCA1. J. Exp. Med. 211, 1027–1036 (2014).

65. Adamo, A. et al. Preventing Nonhomologous End Joining Suppresses DNA Repair Defects of Fanconi Anemia. Mol. Cell 39, 25–35 (2010).

66. Patel, A. G., Sarkaria, J. N. & Kaufmann, S. H. Nonhomologous end joining drives poly(ADP-ribose) polymerase (PARP) inhibitor lethality in homologous recombination-deficient cells. Proc. Natl. Acad. Sci. U. S. A. 108, 3406–3411 (2011).

67. Vidal-Eychenié, S., Décaillet, C., Basbous, J. & Constantinou, A. DNA structure-specific priming of ATR activation by DNA-PKcs. J. Cell Biol. 202, 421–429 (2013).

68. Forment, J. V., Blasius, M., Guerini, I. & Jackson, S. P. Structure-specific DNA endonuclease mus81/eme1 generates DNA damage caused by chk1 inactivation. PLoS One 6, (2011).

69. Ragland, R. L. et al. RNF4 and PLK1 are required for replication fork collapse in ATR-deficient cells. Genes Dev. 27, 2259–2273 (2013).

70. Bertoli, C., Herlihy, A. E., Pennycook, B. R., Kriston-Vizi, J. & De Bruin, R. A. M. Sustained E2F-Dependent Transcription Is a Key Mechanism to Prevent Replication-Stress-Induced DNA Damage. Cell Rep. 15, 1412–1422 (2016).

71. Quinet, A. et al. PRIMPOL-Mediated Adaptive Response Suppresses Replication Fork Reversal in BRCA-Deficient Cells. Mol. Cell 1–14 (2020) doi:10.1016/j.molcel.2019.10.008.

72. Rouaud, F. et al. E2F1 inhibition mediates cell death of metastatic melanoma article. Cell Death Dis. 9, 1–12 (2018).

73. Kakarougkas, A. et al. Opposing roles for 53BP1 during homologous recombination. Nucleic Acids Res. 41, 9719–9731 (2013).

