## Supplemental Table 1 for "ATR Inhibitors as Potent Modulators of DNA End Resection Capacity"

| **Name** | **Sequence (5’-3’)** | **Source** |
| --- | --- | --- |
| NT.1F | CACCGCCAGGCTGAAGTTCGTACCT | This study |
| NT.1R | AAACAGGTACGAACTTCAGCCTGGC | This study |
| NT.2F | CACCGGACCTTCATTGAAGAAAAGC | This study |
| NT.2R | AAACGCTTTTCTTCAATGAAGGTCC | This study |
| NT.3F | CACCGGCTCCCATCCATAGTAAAAA | This study |
| NT.3R | AAACTTTTTACTATGGATGGGAGCC | This study |
| NT.4F | CACCGGGCTTACGTGGGGGGCAAAA | This study |
| NT.4R | AAACTTTTGCCCCCCACGTAAGCCC | This study |
| NT.5F | CACCGACTAGCCTGTTCGCGAGTAG | This study |
| NT.5R | AAACCTACTCGCGAACAGGCTAGTC | This study |
| E2F1.1F | CACCGCGCCGCCGTTGTTCCCGTCA | This study |
| E2F1.1R | AAACTGACGGGAACAACGGCGGCGC | This study |
| E2F1.2F | CACCGGGCCGTGACGGGAACAACGG | This study |
| E2F1.2R | AAACCCGTTGTTCCCGTCACGGCCC | This study |
| E2F1.3F | CACCGTCCGGCGCGTTAAAGCCAAT | This study |
| E2F1.3R | AAACATTGGCTTTAACGCGCCGGAC | This study |
| E2F1.4F | CACCGAAGGATTTGGCGCGTAAAAG | This study |
| E2F1.4R | AAACCTTTTACGCGCCAAATCCTTC | This study |
| E2F1.5F | CACCGTGTCCGGATGGTACCAGGCG | This study |
| E2F1.5R | AAACCGCCTGGTACCATCCGGACAC | This study |
